# Caught a chill? The relationship between canker disease susceptibility and the vulnerability to freeze events in apricot trees

**DOI:** 10.64898/2026.07.12.738016

**Authors:** Guillaume Charrier, Katline Charra-Vaskou, Nathalie Courthieu, Jalayan Lalji, Lia Lamacque, Cindy Morris, Pierre Sudre, Jean-Stéphane Venisse, Christophe Chamet

## Abstract

Bacterial canker remains a major constraint affecting apricot production in South-East of France. It is primarily caused by *Pseudomonas syringae*, a Gram-negative bacterium, many strains of which exhibit ice nucleation activity. By promoting ice formation at relatively high subzero temperatures, ice nucleation-active bacteria may facilitate tissue disruption and pathogen entry. Concurrently, climate-driven shifts toward warmer winter–spring periods have advanced flowering phenology, increasing exposure to late frost events. Despite breeders having developed less susceptible varieties to bacterial canker and early flowering varieties, the link between these traits and frost sensitivity, an emerging risk in this location, remains unresolved. Here, we have evaluated the links between canker susceptibility and frost sensitivity using three cultivar pairs contrasting in disease response and flowering time. Ice nucleation temperature was measured in excised buds under controlled conditions throughout the frost-risk period, alongside field-based diameter variation monitoring over two years. Disease susceptibility (*P* < 0.001), phenology (*P* = 0.003), varieties (*P* < 0.001), locations (*P* < 0.001), and sampling date (*P* < 0.001) significantly affected nucleation temperature, whereas epiphytic bacterial abundance and xylem vessel diameter did not. Trees froze at higher temperatures *in situ* than in laboratory assays (1 to -2°C *versus* -3 to 4°C, respectively), indicating strong environmental modulation of freezing processes beyond Psy-like bacterial activity, which is reflected in contrasting disease susceptibilities (*P* < 0.001) and precocities (*P* < 0.001). These results shed light on the complexity of the freezing process in trees under natural conditions. We discuss the potential roles of microclimatic conditions and alternative ice nucleation sources beyond Psy-like bacteria in driving these physiological processes.

## Introduction

In recent decades, global change has intensified the threat to biological systems, with increasing climate instability and a higher frequency of extreme weather events (Murray & Ebly, 2012) Globalization and rising temperatures have also expanded the ecological niche for plant diseases (Chaloner *et al*., 2021). Among the emerging risks, frost risk lies at the cross-road between biotic and abiotic factors. On the one hand, warming in late winter accelerates the ontogenic development of buds, exposing vulnerable flushing buds to damaging temperatures in spring (Rodrigo, 2000; Charrier *et al*., 2018). Fruit trees are particularly vulnerable to late frosts, with a single freezing event having the potential to destroy the entire year’s production (Melo-Abreu *et al*., 2024). Among fruit trees, *Prunus* species exhibit a wide range of flowering dates, with almonds (*Prunus dulcis*) and apricots (*Prunus armeniaca*) flowering the earliest. On the other hand, in biological tissues, water can remain in a metastable liquid state at temperatures below 0°C (*i*.*e*., supercooling), until ice formation begins around an initial *nucleus*. This process can be catalyzed by ice-nucleating agents, from various origins, including mineral and biological materials (Murray *et al*., 2012). In plants, intrinsic ice nuclei can initiate freezing in the wood of temperate plants (Lamacque *et al*., 2026). Among them, lignin and cellulose (Bogler and Borduas-Dedekind, 2020; Hiranuma *et al*., 2015) have been characterized. External agents are also involved in ice formation, the most famous being the ice nucleation-active bacterium *Pseudomonas syringae* (Morris *et al*., 2013; Lindow, 2023).

In addition to playing a potential role in the freezing of fruit tree tissue, *Pseudomonas syringae* (Psy) can also influence tree health as a plant pathogen. It is the causal agent of many diseases in crops and causes major economic losses in agriculture due to bacterial infection (Mansfield *et al*., 2012). One such disease is bacterial canker, which affects the genus *Prunus* (English *et al*., 1980; Kennelly *et al*., 2007; Vicente *et al*., 2004). Bacterial dieback or bacterial canker caused by Psy has been observed in apricot trees in many areas worldwide, including Bulgaria (Ivanova, 2009), Iran (Karimi-Kurdistani and Harighi, 2008), Italy (Scortichini, 2006) and Turkey (Kotan and Sahin, 2002). Psy belongs to the class of Gammaproteobacteria, is strictly aerobic, Gram-negative, rod-shaped, and motile due to the presence of one to three polar flagella (Young, 2010). Psy is both an endophyte and an epiphyte that colonizes apoplastic tissues, stomata, and vascular systems (Lamichhane *et al*., 2014; Xin *et al*., 2018). Its epidemiological cycle involves dynamic exchanges between internal and surface populations, with peak bacterial growth occurring in spring during leaf emergence and in autumn. The bacteria overwinter in dormant buds and cankers (Hattingh *et al*., 1989; Sundin *et al*., 1988). Psy can enter through natural openings (*e*.*g*., stomata, lenticels, and leaf scars) and wounds caused by biotic factors (pruning and insects) or abiotic factors (frost). Frost-induced lesions provide direct access to the vascular system (Crosse & Garrett, 1966). In apricot trees, the primary natural entry point appears to be the buds (Crosse, 1953; Bordjiba & Prunier, 1989). Therefore, the tree’s phenological stages and anatomical characteristics appear to play a significant role in bacterial population dynamics and, consequently, in predisposition to canker and vulnerability to frost.

For several fruit tree species (peach, apricot, and pear), the freezing potential, in conjunction with tissue freezing temperatures, has been identified as the primary factor for infection during the winter (Weaver, 1978; Klement *et al*., 1984; Montesinos and Vilardell, 1991). The interaction between frost risk and bacterial canker is therefore complex and synergistic. Warming in late winter accelerates bud development, exposing vulnerable tissues to damaging temperatures and creating wounds (Wenneker *et al*., 2012). These mechanical breaks create ideal entry points for the bacteria, which feed on nutrients (*e*.*g*., sugars and amino acids) released by the damaged cells to support growth. In the presence of ice nucleation active bacteria, the freezing point of water increases, enabling parenchyma and cortical tissues to freeze at temperatures approximately 2–3°C higher than the usual values of around -7°C (Lindow, 1987; Morris *et al*., 2013; Lamichhane *et al*., 2014). This reduces the effectiveness of the frost avoidance strategy (Charrier *et al*., 2015; Lamacque *et al*., in press). Frost events create entry points for bacterial infection, while Psy’s ice nucleation activity exacerbates frost damage, creating a feedback loop known as the “frost-bacteriosis complex” (Gaignard & Luisetti, 1993; Klement *et al*., 1974).

Beyond the intensity of the cold, the frequency and number of successive freeze-thaw cycles and the phenological stage of the bud are likely related to the number and severity of infections in apricot trees (Prunier & Bordjiba, 1989; Parisi *et al*., 2019). Dormant plants are particularly susceptible to developing symptoms of bacterial canker (Wilson, 1939; Davis & English, 1969). They become especially sensitive when they remain hydrated during this period (Vigouroux, 1991; Vigouroux & Bussi, 1997). More severe symptoms were recorded when inoculations were performed in midwinter (Garrett, 1979; Endert & Ritchie, 1984). Inoculation and wounding of peach trees in February resulted in more severe canker symptoms than in October and were associated with the necrosis of fruit and shoot buds, as well as delayed budbreak.

Bacterial canker can manifest locally, within the parenchymatous tissues, or systemically, affecting the vascular system and intercellular spaces, resulting in the entry and movement of bacteria through the conducting tissues (Lamichhane *et al*., 2014; Lindow, 2023). However, *P. syringae*-induced vascular diseases have been observed in deciduous but not in evergreen trees (Ramos & Kamidi, 1981; Young, 2004; Green *et al*., 2009; Hattingh *et al*., 1989; Kennelly *et al*., 2007; Lamichhane *et al*., 2014).

*P. syringae* can be found in the interior tissues of fruiting cherry trees, including the branches and trunk (Cameron, 1970). This leads to significant degradation and browning along the length of the vessels from the infected area, which can result in the death of the tree (Omrani, 2018). The relationship between ice formation and bacterial infection highlights the importance of the vascular structure of plants in their ability to withstand abiotic and biotic stresses. Ice formation and the spread of bacteria directly impact the functioning of the vascular system, which is essential for trees.

In the last decade, apricot production in Southeast of France has been heavily affected by bacterial canker and spring freezes. Although breeders have developed less susceptible varieties to bacterial canker and early flowering varieties, the link between these traits and frost sensitivity, an emerging risk in this location, is still obscure. In this study, we investigated the relationship between the susceptibility to bacterial canker, phenological precocity, wood anatomy, and ice nucleation temperature in various apricot varieties. We hypothesized that varieties with a higher susceptibility to canker would be able to host a larger population of ice nucleation active bacteria, resulting in a higher ice nucleation temperature in their tissues than if they only had internal ice nuclei. Furthermore, we hypothesized that susceptible varieties would have larger xylem conduits to allow pathogen propagation and that early-blooming varieties would exhibit even higher sensitivity to freeze-thaw cycles. To test these hypotheses, we measured the freezing threshold temperature under controlled and field conditions, across three pairs of varieties with contrasting phenology (early, intermediate, and late flowering) and susceptibility to bacterial canker. Additionally, we measured Psy-like bacterial abundance on buds and examined the anatomy of the xylem vessels.

## Material and Methods

### Ice nucleation temperature in excised stems

#### Plant material

Apricot trees (*Prunus armeniaca* L.) were sampled in experimental orchards at two locations in South-East of France, in Etoile sur Rhone (SEFRA: N 44.817° E 4.884°) and Toreilles (Centrex: N 42.756° E 2.982°) between February and April 2023. In both orchards, six varieties were selected according to their phenology (early, intermediate and late) and tolerance to bacterial canker (susceptible *vs* less susceptible; Tab. 1). Every two weeks, between February 2^nd^ and April 2^nd^, two to three twigs bearing 6 to 10 buds each were sampled in SEFRA and CENTREX orchards. Samples were shipped within 24 to 48 hours to the laboratory for exotherm analysis. The phenological stages of the reproductive organs studied ranged from dormant buds (BBCH 50) to small fruits (BBCH 77).

#### Exotherm analysis

The samples (approx. 10 cm long), bearing one to three individual buds were inserted into wet rock wool and placed in a vertical position. One thermocouple (type T) was inserted under the surface scales of each bud (BBCH 50 to 57), inside the carpel (BBCH 59 to 69) or inside the fruit (BBCH 70 to 77), depending on the phenological stage. All thermocouples (n = 56) were connected to a data logger with two multiplexers (CR1000 and AM25T, Campbell, Logan, Utah, USA). Temperature was recorded three times per minute and averaged every minute.

After temperature stabilization at 5°C for 30 minutes, samples were exposed to a standardized freeze-thaw cycle down to -25°C at a rate of 2.5 K.h^-1^ in a freezing chamber (Model MK, Binder, Germany). During the freezing stage, the ice formation exotherm results in a rapid temperature rise of around 1 to 3°C, which dissipates in about ten minutes. The temperature of the organ at which the exothermic reaction is detected was considered as the ice formation temperature (Ice Nucleation Temperature, INT; °C).

#### Microbiology

The abundance of *Pseudomonas syringae*-like (Psy-like) bacteria on buds was determined in samples from SEFRA and Centrex in March 2025. Three buds from the same branch were immersed in 10mL of NaCl (0.8% w/v) and shaken overnight at 4°C. The suspension containing epiphytic bacteria was dilution-plated on King’s medium B (KB) plates (King *et al*., 1954) and incubated for 48 h at 25°C before counting the colonies. Colonies were observed under UV light to detect pyoverdine fluorescence in colonies exhibiting typical Psy-like morphology colonies and counted. Population sizes of Psy-like bacteria were calculated as colony-forming units (CFU) normalized by total bud fresh weight. When no CFUs were observed growing on the plate, they were considered below the detection threshold and assigned the value of 100 CFU.bud^-1^; Hirano *et al*., 1982).

#### Anatomy

Three shoots, each approximately 50 cm long, were collected from three individual of each variety for the study of vascular anatomy. After sampling, the shoots were stripped of their leaves and stored at 4°C under humid conditions until preparation. Transverse sections were made from a stem segment located approximately 45 cm from the apex, at the middle of an internode. These samples were immersed for 30 minutes in a 15:85 Ethanol:water solution under vacuum.

The anatomical sections were prepared on an RM2165 microtome (Leica, Germany) coupled with a Julabo 600F cooling system and a Basetech BT-305 electrical system to secure the sample using the Peltier effect. Sections 25 µm thick were prepared along the longitudinal axis of the stem. The sections were then stained with 0.5% (w/v) toluidine blue, rinsed with water and mounted between a slide and a cover slip in a 1:1 water-glycerol mixture.

Observations of the cross-sections were carried out using an Axio Zoom.V16 macroscope to obtain a general view of the section, followed by a Zeiss Axioplan 2 optical microscope to examine the vessels in greater detail. The diameter, density and surface area of each vessel were measured in the stems of each variety using ImageJ software.

### Detection of ice nucleation in the field

To detect ice formation in natural conditions, automated high-resolution dendrometers and temperature loggers (PepiPIAF, CaptConnect, France) were installed on one north-exposed main branch on three individual trees from six varieties, in the experimental orchard of SEFRA (March 1^st^ 2023). These varieties were selected according to their phenology (early, intermediate and late) and tolerance to bacterial canker (susceptible *vs* less susceptible). Branch diameter and logger internal temperature were measured every 30 minutes. When ice forms, a rapid and large shrinkage (*e*.*g*., several hundreds of µm within one hour) is detected, due to ice-induced dehydration of bark living cells (Charra-Vaskou *et al*., 2016; Charrier *et al*., 2017). The temperature measured with the datalogger at the moment of shrinkage was considered as the ice nucleation temperature (INT; °C).

### Statistics

Statistical differences between ice nucleation temperature of different groups (varieties, dates, location and covariables) were tested according to Kruskal-Wallis non-parametric test, as the data, in most cases, were not normally distributed according to Shapiro-Wilk test.

Ice nucleation temperature in excised stems was predicted by generalized linear models using relevant quantitative variables (glm function in R ver 4.3.1). Among the variables, the cumulative chilling hours, with mean temperature between 0 and 7.16°C, from September 1^st^ and cumulative growing degree days, with base temperature at 0°C, from January 1^st^ were computed from local weather stations. The best model was selected based on the accuracy of the prediction *via* the Root Mean Standard Error (RMSE) and parsimony, *via* the Bayesian Information Criterion (BIC).

In the field, ice nucleation temperature across varieties was fitted using a logistic function with temperature as the explanatory variable (nls function in R ver 4.3.1).

## Results

### Experimental freezing assays

The ice nucleation temperature (INT) in excised stems varied within a range of two degrees, from - 2.30 ± 0.05°C (Rubissia in early March) to -4.36 ± 0.08°C (Farbela in mid-February). Significant differences were observed according to the susceptibility to bacterial canker with higher INT values in susceptible varieties (median = -3.69 *vs* -3.38°C in less susceptible *vs* susceptible varieties, respectively; P < 0.001; Fig. 1A). Significant differences were also observed according to phenological stage (P < 0.001; Fig. 1B). However, the trend of higher INT with phenological development was not monotonic. Specifically, full bloom (BBCH 67), late bloom (BBCH 69) and large fruits (BBCH 77) exhibited INT that were 0.5 to 0.8°C higher than the rest. Early (*i*.*e*., from BBCH 50 to 61) and late (*i*.*e*., between BBCH 71 and 75) phenological stages were not significantly different. In terms of precocities, early and late varieties had a lower INT than intermediate ones (P = 0.003; Fig.1C). Among the other studied factors, significant differences were observed across varieties (P < 0.001; Fig.1D), locations (P < 0.001; Fig.1E) and dates (P < 0.001; Fig.1F).

**Figure 1.**
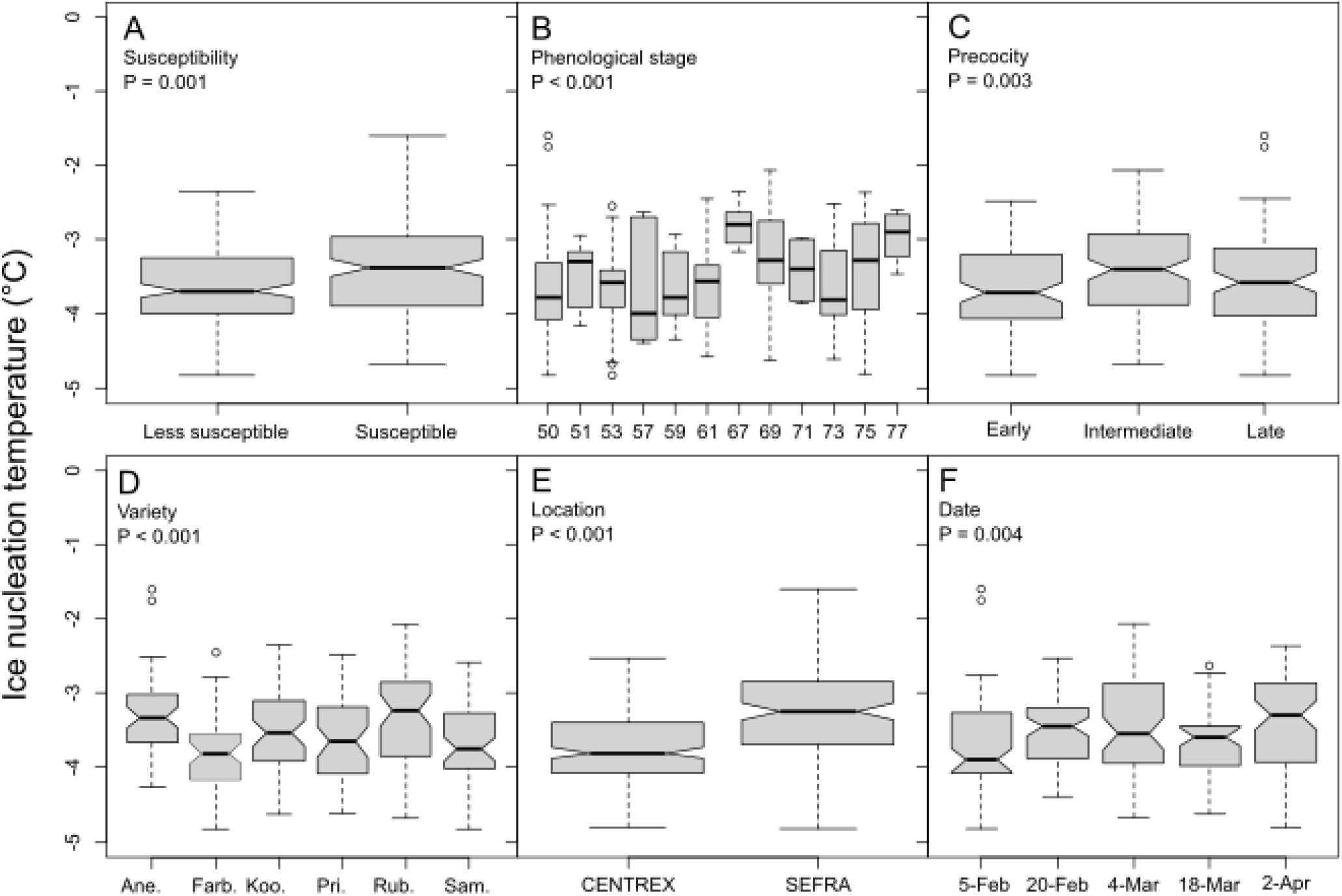
Ice nucleation temperature in apricot flower buds, as measured in controlled chamber experiments according to various factors: susceptibility to bacterial canker (Susceptibility; A), phenological stage (B), Precocity (C), variety (D), location (E) and sampling date (F). The *p*-value according to the Kruskal–Wallis non-parametric test indicates whether the factor has a statistically significant effect. Empty circles indicate outliers.

The abundance of epiphytic Psy-like bacteria on buds (BBCH 59) of these varieties, was highly variable (around 10^4^ CFU per bud; Fig. 2). While a few buds hosted extremely high numbers of bacteria (> 10^5^ CFU per bud), 11% of buds harbored only a few hundreds of bacteria per bud. Although significant differences in INT were observed across varieties and susceptibility to bacterial canker, the abundance of epiphytic Psy-like bacteria was not significantly related to susceptibility to bacterial canker (P = 0.755), or variety (P = 0.497). As a consequence, no relation with INT was observed (R^2^ = 0.089; P = 0.347).

**Figure 2.**
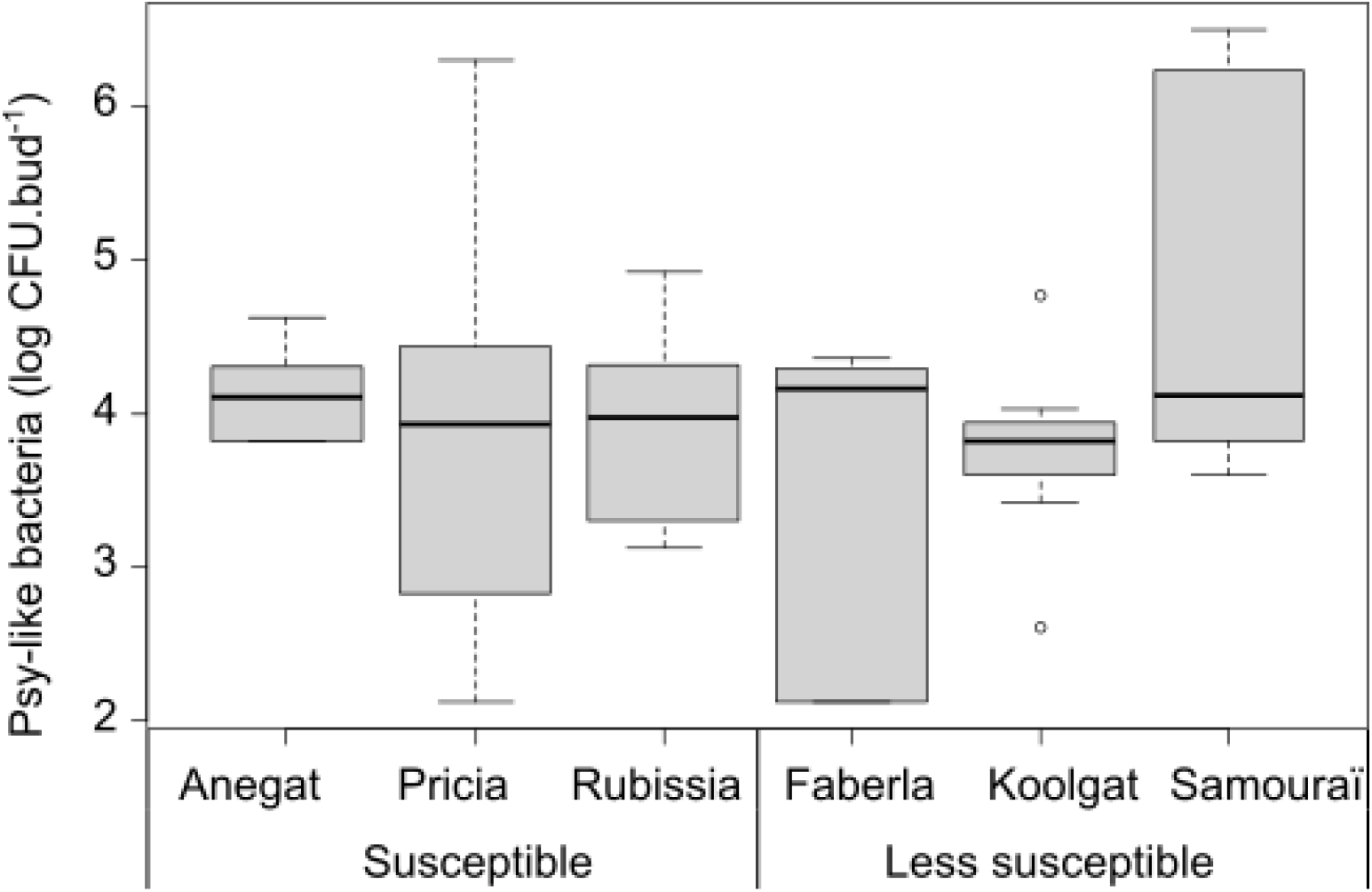
Abundance of Psy-like bacteria on buds of different varieties of apricot. Stems of different apricot varieties had similar vessel diameter (approx. 20-25 µm) with no significant differences across varieties (P = 0.939) or susceptibility to bacterial canker (P = 0.847; Fig. 3). Theoretical hydraulic conductance (approx. 0.04-0.05 g.m-1.MPa-1.s-1) was also highly similar with respect to varieties (P = 0.519) or susceptibility to bacterial canker (P = 0.700).

**Figure 3.**
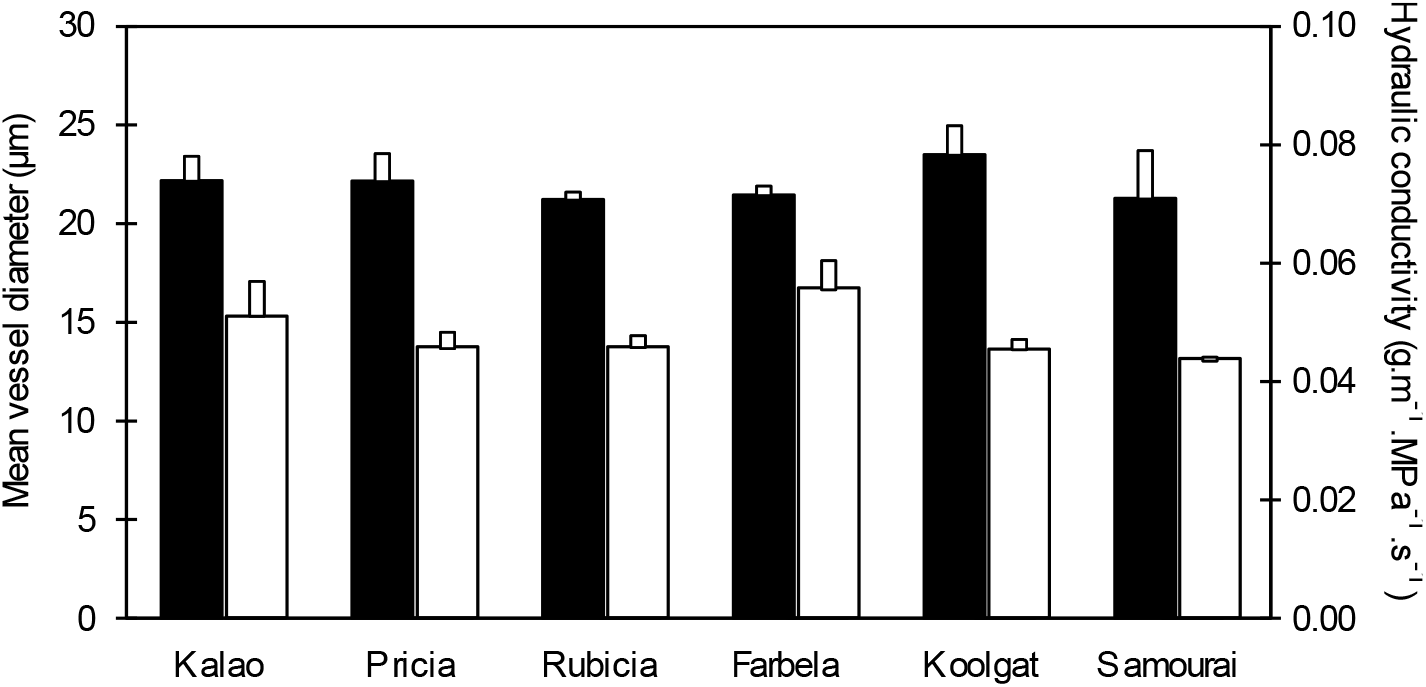
Mean vessel diameter (solid bar) and theoretical hydraulic conductivity (empty bar) in different varieties of apricot (mean + SE).

Of the variables that were significantly correlated with INT, all were also correlated with phenological stage, either directly (*e*.*g*., precocity and variety) or indirectly (*e*.*g*., location and date). We examined the distinct effects of cumulative chilling and forcing temperatures on INT. Cumulative chilling hours from September were tightly correlated with INT (R^2^ = 0.182; P < 0.001). Conversely, cumulative forcing hours from January were only slightly correlated with INT (R^2^ = 0.009; P = 0.090). Finally, the best additive linear model based on BIC predicts INT using susceptibility to bacterial canker, sampling date, and cumulated forcing units at the sampling date as the relevant variables (R^2^ = 0.241; BIC = 510.6). This model accurately predicts the freezing temperature (RMSE = 0.525°C; Figure 4).

**Figure 4.**
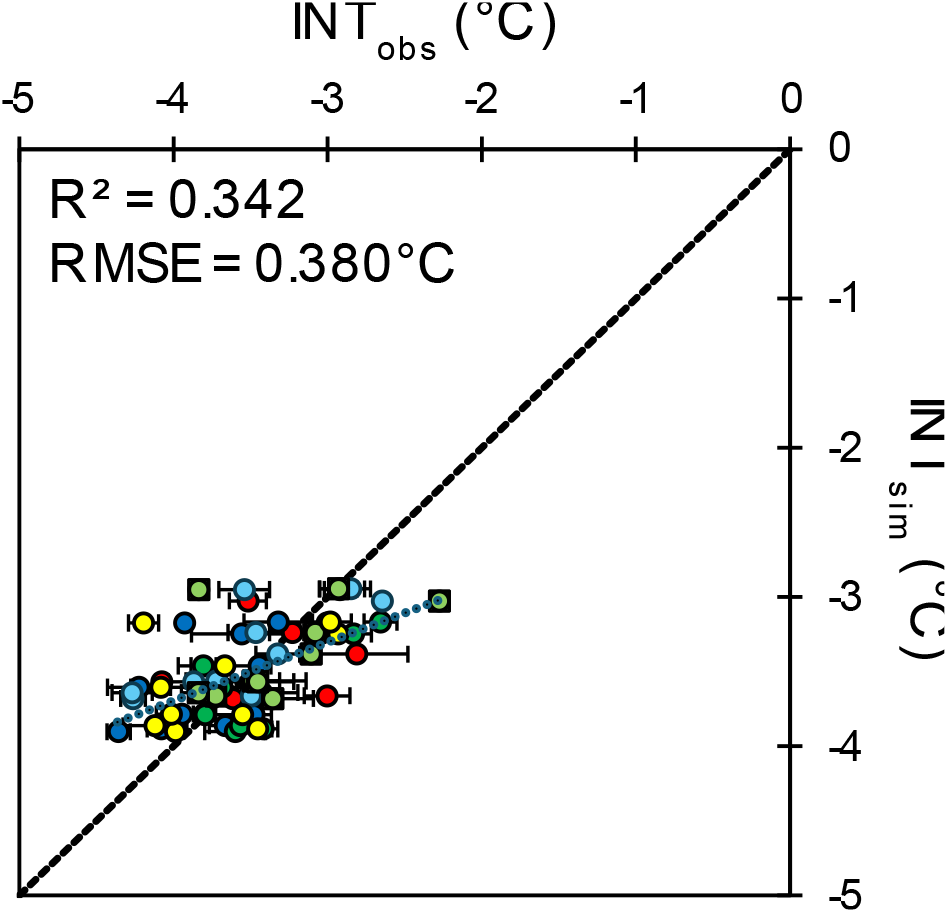
Predicted Ice Nucleation Temperature (INT_sim_) based on observed Ice Nucleation Temperature (INT_obs_). The solid line corresponds to INT_sim_ = INT_obs_.

#### Field measurement of INT

During the spring of 2023, temperatures remained almost constantly above 0°C. While a few meteorological freezes were recorded depending on the local conditions of each tree (daily minimum temperature < 0°C), these events triggered only three physiological freezes, detected by a rapid shrinkage of the branch upon freezing, in the monitored trees. During the winter of 2023–2024, the individual trees experienced between 18 and 24 meteorological freezes (T_min_ < 0°C), which resulted in between 12 and 16 physiological freezes (Table 1). No significant difference was observed in the number of freezes across varieties (P = 0.682), susceptibility to bacterial canker (P = 0.777) or precocity (P = 0.952).

**Table 1.** Number of meteorological freezes (temperature < 0°C), number of physiological freezes (stem diameter shrinkage) and ice nucleation temperature in the branches of different apricot varieties in the SEFRA orchard.

| Variety | Susceptibility | Precocity | Meteorological freeze | Physiological freeze | $T_{freeze}$ |
| --- | --- | --- | --- | --- | --- |
| Memphis | Low | Early | $24.00 \pm 1.17$ | $15.75 \pm 2.30$ | $-0.48 \pm 0.39$ |
| Koolgat | Low | Medium | $22.33 \pm 4.46$ | $12.67 \pm 0.72$ | $-2.13 \pm 0.54$ |
| Lisa | Low | Late | $21.67 \pm 4.28$ | $13.33 \pm 1.91$ | $-2.03 \pm 0.66$ |
| Pricia | High | Early | $18.00 \pm 1.77$ | $14.50 \pm 1.03$ | $-1.38 \pm 0.81$ |
| Rubissia | High | Medium | $19.33 \pm 1.44$ | $12.33 \pm 0.54$ | $-0.75 \pm 0.49$ |
| Anegat | High | Late | $22.00 \pm 1.27$ | $14.50 \pm 1.92$ | $-1.75 \pm 0.68$ |

However, significant differences in the ice nucleation temperature were observed across varieties (P < 0.001), susceptibility to bacterial canker (P < 0.001), precocity (P < 0.001) and cumulative chilling hours (P < 0.001) but it was not correlated to cumulative growing degree-days (P = 0.087). It should be noted that, in contrast to laboratory measurements, INT is lower in early and late varieties than in intermediate ones. The relationship between the probability of ice formation and temperature varied across varieties with freezing temperatures ranging from -0.48°C (Memphis) to -2.13°C (Koolgat). Although the temperatures observed in the field were higher than those observed under controlled conditions (between -2.30 and -4.36°C), overall, the most susceptible varieties show a slightly higher temperature at which freezing occurred (-1.16 ± 0.14°C and -1.46 ± 0.21°C for susceptible *vs* less susceptible varieties, respectively; Figure 5).

**Figure 5.**
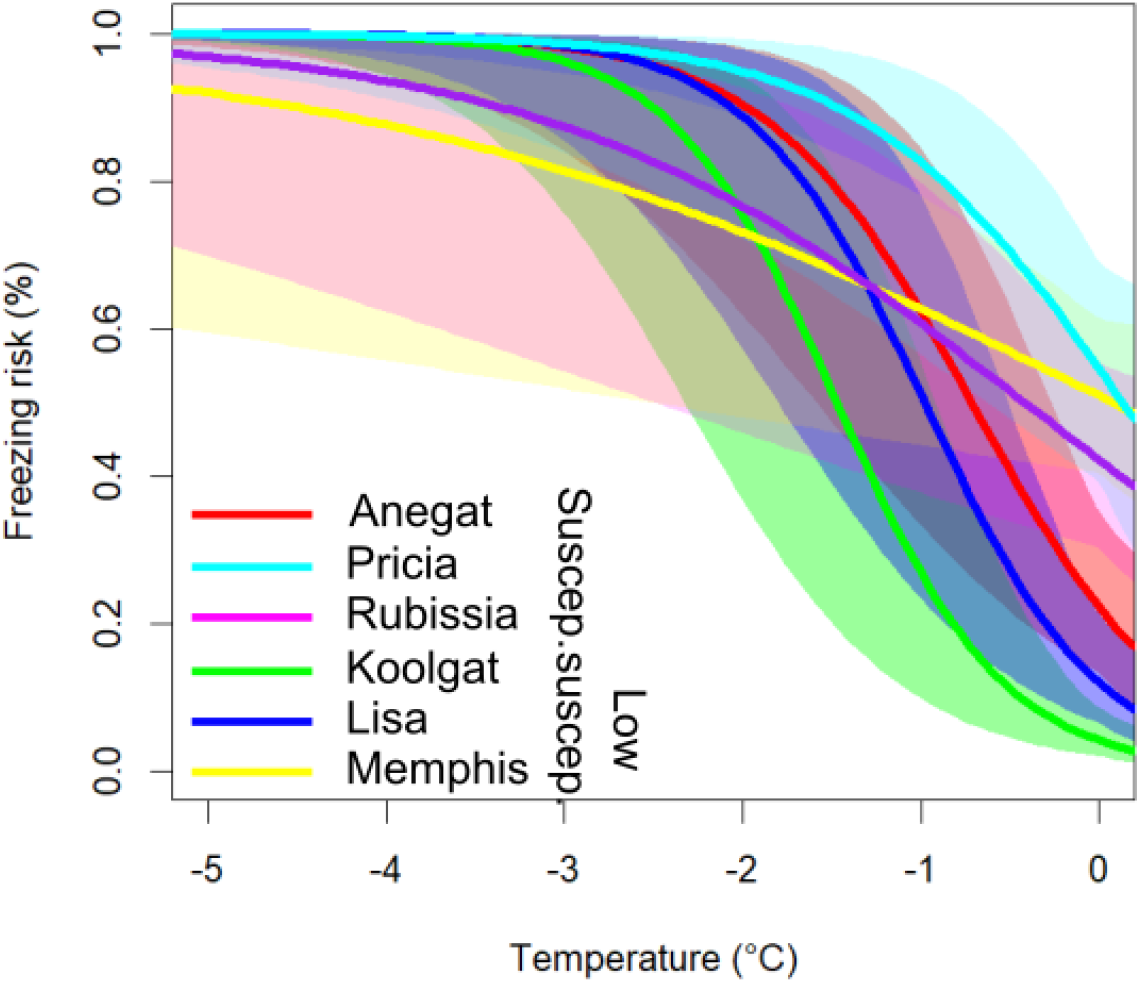
Probability of ice formation (freezing risk) in the field depending on the temperature in the stem of different apricot varieties. Shaded area represents the confidence interval at 90% for the corresponding variety.

## Discussion

Significant differences in ice nucleation temperature were observed across apricot varieties, in relation to their susceptibility to bacterial canker and their precocity. However, no significant relationship was observed with the abundance of Psy-like bacteria, suggesting that varieties with a higher susceptibility to bacterial canker do not consistently harbor larger populations of *P. syringae* on their buds at the times when freezing risk occurs. Moreover, the absence of a relationship with vessel diameter suggests that the propagation of pathogens through xylem tissue does not depend primarily on vessel diameter.

As hypothesized, we showed that varieties most susceptible to bacterial canker had a higher freezing point, rendering them more susceptible to freezing injury than the least susceptible varieties. Although the difference in the mean ice nucleation temperature was approximately 0.3°C (1.5°C when comparing the most extreme varieties) in both controlled conditions and the field, the difference was significant, resulting in approximately two additional freeze-thaw cycles experienced by the trees in the field each year. Exposure to an increased number of freeze-thaw cycles can have a detrimental effect on xylem hydraulic conductivity (Charrier *et al*., 2014), particularly when apricot trees are unable to refill embolized vessels in spring (Améglio *et al*., 2002). However, freeze-thaw cycles did not significantly stimulate longitudinal water flux in the xylem tissues (data not shown). At the cellular level, freeze stress is mainly associated with dehydration, membrane destabilization, and reactive oxygen species generation, requiring the accumulation of osmoprotectants and the activation of antioxidant machinery, which can influence freezing tolerance (Theocharis *et al*., 2012; Charrier *et al*., 2013; Ding *et al*., 2019).

Surprisingly, despite the absence of a clear link between Psy-like bacteria abundance and INT, susceptible varieties were more prone to freezing, with higher INT. This phenomenon was previously reported in five fruit tree species where no significant relationships between ice nucleation-active bacteria abundance, INT, and frost damage were reported (Proebsting & Gross, 1988). This may be due to a reduced ice nucleation activity in the Psy-like bacteria present in our samples. Ice nucleation activity in *P. syringae* is controlled by outer membrane Ina proteins encoded by the *inaZ* gene, whose expression and aggregation state directly determine ice nucleation efficiency (Green & Warren, 1985; Govindarajan & Lindow, 1988). Variability in *ina* gene expression or post-translational assembly, along with the accumulation of various cryoprotectants (*e*.*g*., glycine betaine, exopolysaccharides, Cold-shock proteins, and Antifreeze proteins) could therefore explain discrepancies between bacterial abundance and effective ice nucleation activity (Phadtare, 2004 ; Li *et al*., 2013 ; Lorv *et al*., 2014 ; Carrion et al, 2015). In the field, such a manipulation has been achieved by spraying non-ice nucleation-active strains of Psy-like bacteria, obtained through mutation of the *ina* gene, to reduce the ratio of ice nucleation-active strains on pear trees and, consequently, reduce frost damage (Cody *et al*., 1987; Lindow, 1987). The abundance of epiphytic ice nucleation-active bacteria is probably not as critical, with internal ice *nuclei* playing a more significant role. Dynamic interactions between the internal and epiphytic populations can mask a significant proportion of ice nucleation-active bacteria (Hattingh *et al*., 1989). Phloem tissue is indeed a significant reservoir for Psy populations, which increase when exposed to cold temperatures (Klement *et al*., 1984). Cold exposure can also induce the expression of virulence-associated genes, including those involved in the type III secretion system (T3SS), enhancing bacterial fitness and host colonization under stress conditions (Xin *et al*., 2018). Although it is usually recommended to avoid pruning during the freezing period to limit bacterial development, we did not observe significant differences in xylem anatomy between susceptible and less susceptible varieties. Wood structure is indeed relatively similar and should not affect the ability of the pathogen to propagate in the wood.

Another possibility is that the relationship between the abundance of ice nucleation-active bacteria and INT is indirect, plausibly mediated by the host’s systemic immune responses (Kour *et al*., 2024), and that Psy values measured in this study are not a direct measure of the exposure to the bacterial canker pathogen, as not all Psy cause disease on apricot. The immune response to Psy may rely on distinct mechanisms, such as stopping infection and limiting progression (Omrani, 2018), which could explain why, although susceptibility to bacterial canker is a significant factor, the effect on INT varies across varieties, with Koolgat exhibiting a higher INT in the laboratory (Fig. 1) and Memphis a higher INT in the field (Table 2). Further studies should detail the specific disease resistance mechanisms in these varieties.

**Table 2.** Spearman’s rho coefficients across varieties, susceptibility levels, and precocities for INT in the laboratory and in the field, Psy-like abundance (Psy), vessel diameter (D), hydraulic conductance (K_H_), and the number of freezing events experienced in the field (FE_Field_)

|  | Variety | Susceptibility | Precocity |
| --- | --- | --- | --- |
| $INT_{\text{lab}}$ | <0.0001 | 0.0035 | 0.0010 |
| $INT_{\text{field}}$ | <0.0001 | <0.0001 | <0.0001 |
| Psy | 0.4971 | 0.4589 | 0.7555 |
| D | 0.9388 | 0.8809 | 0.8474 |
| $K_H$ | 0.5188 | 0.3102 | 0.7003 |
| $FE_{\text{Field}}$ | 0.6824 | 0.9523 | 0.7767 |

Finally, variability in INT and frost damage can result from other intrinsic nucleators besides ice nucleation-active bacteria (Lamacque *et al*., 2026). In studies relating ice nucleation-active bacteria and INT, the relationship was not always significant in grapes (Itier *et al*., 1991) and other tree species (Proebsting & Gross, 1988). It has been suggested, for example, that ice nucleation-active bacteria can contribute to INT as low as -2°C, whereas non-bacterial ice nucleation agents contribute to INT at around -4°C (Gross *et al*., 1988). In plants, intrinsic ice nucleators may include cell wall components, proteins, or lipid bodies, whose physicochemical properties and spatial organization influence ice initiation (Pearce, 2001; Bredow & Walker, 2017; Debnath *et al*., 2026).

In the field, the INT was consistently higher than in controlled conditions, on excised stems. This difference has been observed previously (Ashworth & Davis, 1987; Proebsting & Gross, 1988) and can be explained by the stochasticity of the ice nucleation process. Lower INTs are observed in excised buds than in buds that are still attached to a piece of stem (Proebsting & Gross, 1988). In our study, although the buds remained attached, the number of ice nucleation points was likely lower, as this number decreases with the sample size, thereby reducing the probability of freezing at a given temperature. With a higher number of nucleation points, ice can propagate over a greater distance in supercooled woody tissues in the field (Charrier *et al*., 2017). In peach trees, the relationship between INT and sample weight stabilizes at approximately 5 g of fresh weight (Ashworth & Davies, 1984). Our samples consistently exceeded this threshold and were in the plateau of the logarithmic relationship between INT and the sample’s fresh weight (Gross *et al*., 1988). Finally, as only a single nucleus is necessary to trigger freezing, INT in the field occurs at much higher temperatures, resulting in small but significant differences in the freezing behavior of up to 1.5°C across the most extreme apricot varieties.

Phenological precocity and actual phenological stage were also significant factors explaining INT dynamics. During spring bud ontogenic development, tissue water content increases significantly reaching 65% or more of the total mass (Charrier *et al*., 2013). Water content is a significant factor explaining INT in the xylem (Lintunen *et al*., 2013), thus influencing bud INT when the hydraulic connection between stem and bud is restored (Vilouta *et al*., 2022). At the molecular level, this transition is associated with the regulation of various integral membrane channels, such as aquaporins (*e*.*g*., plasma membrane intrinsic proteins, PIPs), and cell wall remodeling enzymes, which modulate water transport and tissue permeability during de-acclimation (Rahman *et al*., 2020). The resumption of hydraulic connection becomes key in the springtime when buds change their cold resistance strategy from cold tolerance to cold avoidance (Charrier *et al*., 2015). However, the reversal of the ranking with respect to precocities between field monitoring and laboratory experiments suggests a complex interaction between freezing patterns and growth resumption, which does not directly translate from the organ scale to the whole tree scale, where structural heterogeneity buffers temperature changes and water relations.

## Conclusion

In conclusion, we showed that the freezing behavior of different apricot varieties differs substantially in relation to their susceptibility to bacterial canker, while no significant differences were observed in the abundance of Psy-like bacteria on bud surfaces. These findings suggest that the greater frost vulnerability observed in susceptible varieties is not directly driven by epiphytic populations of *P. syringae*, but is more likely associated with other ice-nucleating agents or intrinsic plant properties. In particular, structural and physiological traits, as well as internal microbial populations, may play a key role in modulating ice nucleation and propagation within tissues. More broadly, our results highlight that frost sensitivity arises from a complex interplay between biotic and abiotic factors.

Further studies should therefore integrate additional drivers of freezing and disease dynamics, such as soil texture, low soil pH, soil depth, tree nutrition, tree age, rain, rootstock selection, height of grafting, and early autumn pruning, as well as orchard microclimatic conditions. Likewise, at the molecular level, both plant regulatory mechanisms involved in cold acclimation and bacterial determinants of ice nucleation activity deserve further investigation to better understand their combined effects on frost damage. Finally, the contribution of the *P. syringae* ice nucleation-active population to frost injury is likely limited, as ice nucleation *in planta* is also governed by intrinsic nucleators and environmental conditions, rather than being solely dependent on bacterial INA activity.

## Aknowledgements

We are very thankful to Thierry Améglio, Laurent Barroux, and Pascal Walser for their assistance with field monitoring and sampling. We would also like to thank Pierre Amato and Nicolas Rouchon for their help with laboratory measurements. This project was funded by France-Agri-Mer through the CASDAR “*Connaissance”* program (Vaccin project).

## Notes

### Competing Interest Statement

The authors have declared no competing interest.

